# Restoration of *Sp4* in forebrain GABAergic neurons rescues hypersensitivity to ketamine in *Sp4* hypomorphic mice

**DOI:** 10.1101/015677

**Authors:** Kerin K. Higa, Baohu Ji, Mahalah R. Buell, Victoria B. Risbrough, Susan B. Powell, Jared W Young, Mark A. Geyer, Xianjin Zhou

## Abstract

Ketamine produces schizophrenia-like behavioral phenotypes in healthy people. Prolonged ketamine effects and exacerbation of symptoms were observed in schizophrenia patients after administration of ketamine. More recently, ketamine has been used as a potent antidepressant to treat patients with major depression. The genes and neurons that regulate behavioral responses to ketamine, however, remain poorly understood. Our previous studies found that *Sp4* hypomorphic mice displayed several behavioral phenotypes relevant to psychiatric disorders, consistent with human *SP4* gene associations with schizophrenia, bipolar, and major depression. Among those behavioral phenotypes, hypersensitivity to ketamine-induced hyperlocomotion has been observed in *Sp4* hypomorphic mice. Here, we report differential genetic restoration of *Sp4* expression in forebrain excitatory neurons or GABAergic neurons in *Sp4* hypomorphic mice and the effects of these restorations on different behavioral phenotypes. Restoration of *Sp4* in forebrain excitatory neurons did not rescue deficient sensorimotor gating, fear learning, or ketamine-induced hyperlocomotion. Restoration of *Sp4* in forebrain GABAergic neurons, however, rescued ketamine-induced hyperlocomotion, but did not rescue deficient sensorimotor gating or fear learning. Our studies suggest that the *Sp4* gene in forebrain GABAergic neurons plays an essential role in regulating some behavioral responses to ketamine.

## Introduction

Sp4 is a transcription factor that recognizes GC-rich sequences in the “CpG islands” around the promoters of many genes (Heisler *et al*, 2005). In contrast to the *Sp1* gene, *Sp4* gene expression is restricted to neuronal cells in brain (Supp *et al*, 1996; Zhou *et al*, 2005). Our previous studies found that *Sp4* hypomorphic mice displayed several behavioral phenotypes relevant to psychiatric disorders (Zhou *et al*, 2005; Zhou *et al*, 2010; Zhou *et al*, 2007). In humans, the *SP4* gene was reported to be sporadically deleted in patients with schizophrenia (Tam *et al*, 2010; Zhou *et al*, 2010), and the SP4 protein is reduced in the postmortem brains of bipolar patients (Pinacho *et al*, 2011). Additionally, human genetic studies reported the association of *SP4* gene with bipolar disorder, schizophrenia, and major depression (Greenwood *et al*, 2011; Shi *et al*, 2010; Shyn *et al*, 2009; Zhou *et al*, 2009). These studies suggest the *Sp4* hypomorphic mice as a promising mouse genetic model for human psychiatric disorders.

Administration of ketamine, a noncompetitive NMDAR antagonist, induces behaviors that resemble several aspects of psychiatric disorders in healthy people (Krystal *et al*, 1994). Prolonged ketamine effects and exacerbation of symptoms were reported in schizophrenia patients (Holcomb *et al*, 2005; Lahti *et al*, 1995; Malhotra *et al*, 1997). On the other hand, ketamine was recently found to act as a potent antidepressant that provides rapid relief to patients who are resistant to treatment with classical antidepressants (aan het Rot *et al*, 2010; Diazgranados *et al*, 2010; Price *et al*, 2009; Zarate *et al*, 2006). It is unclear however, what genes and neurons are the key regulators of such responses to ketamine.

*Sp4* hypomorphic mice have robust behavioral phenotypes such as deficient prepulse inhibition (PPI) and fear learning deficits, as well as hypersensitivity to ketamine-induced hyperlocomotion (Ji *et al*, 2013). By taking advantage of a genetic rescue strategy used in the generation of *Sp4* hypomorphic mice (Zhou *et al*, 2005), we restored *Sp4* expression in forebrain excitatory and GABAergic neurons, respectively. Behavioral deficits in PPI and fear conditioning as well as ketamine–induced hyperlocomotion were examined in the two different neuron-specific *Sp4* rescue mice.

## Materials and Methods

### Mouse strains and breeding

The *Sp4* hypomorphic mice were generated as previously described (Ji *et al*, 2013). *Emx1-Cre* mouse line (stock number: 005628) and *Dlx6a-Cre* mouse line (stock number: 008199) were purchased from Jackson laboratory. Both lines were backcrossed with Black Swiss mice for more than 6 generations. Each *Cre* line was then bred with *Sp4* heterozygous mice on Black Swiss background to obtain double heterozygous mice. The double heterozygous mice were finally bred with *Sp4* heterozygous mice on 129S background to obtain F1 generation mice. All F1 *Sp4* heterozygous mice were sacrificed, and the remaining four genotypes were used for molecular and behavioral analyses. Mice were housed in a climate-controlled animal colony with a reversed day/night cycle. Food (Harlan Teklab, Madison, WI) and water were available *ad libitum*, except during behavioral testing. Behavioral testing began at 3 month-old mice with prepulse inhibition, fear learning, and ketamine-induced locomotor activity with at least 2 weeks inbetween tests. All testing procedures were approved by the UCSD Animal Care and Use Committee (permit number: A3033-01) prior to the onset of the experiments. Mice were maintained in American Association for Accreditation of Laboratory Animal Care approved animal facilities at UCSD and local Veteran’s Administration Hospital. These facilities meet all Federal and State requirements for animal care.

### LacZ staining

20 µm thick cryostat sections were cut from fresh adult mouse brains. The sections were fixed, permeabilized, and stained as previously described (Zhou *et al*, 2005).

### RNA and protein expression analyses

RNA was extracted from mouse cortex and striatum from different genotypes using RNeasy Lipid Tissue Mini Kit (Qiagen). RT-PCR was conducted as previously described (Ji *et al*, 2014). Total protein was extracted from mouse cortex and striatum and Western blot analyses were subsequently performed as described (Ji *et al*, 2014). Anti-Sp4 antibody was purchased from Santa Cruz Biotechnology (sc-13019, sc-645).

### PPI and VT locomotion test

PPI and VT locomotion tests were conducted as previously described (Ji *et al*, 2013).

### Ketamine

Ketamine was dissolved in saline and administered i.p. at a volume of 5 ml/kg after 30 minutes of habituation in the VT arena (Brody *et al*, 2003). The doses of ketamine were determined from the same F1 genetic background mouse in previous studies (Ji *et al*, 2013). A within-subjects crossover design was used for drug studies, with 1 week between drug treatments.

### Statistical analysis

Repeated measures analysis of variance (ANOVA) with genotype as a between-subjects factor and drug treatment, block, and prepulse intensity as within-subjects factors were performed on the %PPI data and total distance traveled. *Post hoc* analyses were carried out using Newman-Keuls or Tukey’s test. Alpha level was set to 0.05. All statistical analyses were carried out using the BMDP statistical software (Statistical Solutions Inc., Saugus, MA).

## Results

### Generation of neuron-specific *Sp4* rescue mice

Mouse *Emx1* (Briata *et al*, 1996) and *Dlx5/6* (Robledo *et al*, 2002) expression are first detected at embryonic E9.5 and E8.5, respectively. Forebrain excitatory neurons originate from the *Emx1* cell lineage (Gorski *et al*, 2002), while forebrain GABAergic neurons are derived from *Dlx5/6* cell lineage (Monory *et al*, 2006). Since mouse *Sp4* expression starts around E9.5 during embryogenesis (Supp *et al*, 1996), the *Cre* driven by *Emx1* and *Dlx5/6* genes will restore *Sp4* gene during the entire development of forebrain excitatory and GABAergic neurons, respectively. Both *Emx1-Cre* and *Dlx6a-Cre* mouse lines were purchased from Jackson laboratory and backcrossed with Black Swiss mice for more than 6 generations. Each *Cre* line was further bred with Black Swiss *Sp4* heterozygous mice to generate double heterozygous mice that were finally bred with *Sp4* heterozygous mice on 129S background to generate F1 mice (Figure 1S). In F1 *Sp4* homozygous mice carrying either the *Emx1-Cre* or *Dlx6a-Cre* (*Dlx5/6-Cre*) gene, the nuclear *LacZ* cassette was excised to restore *Sp4* expression in each cell lineage (Figure 1). Consistent with previous reports on *Emx1-Cre* expression (Gorski *et al*, 2002), introduction of *Emx1-Cre* into *Sp4* heterozygous mice abolished *LacZ* staining in both cortical and hippocampal excitatory neurons (Figure 1a). Under higher magnification, many *LacZ* blue spots remained in both cortex and hippocampus. They presumably consisted of GABAergic and other types of neurons (Figure 1b). Introduction of *Dlx6a-Cre* into *Sp4* heterozygous mice abolished *LacZ* staining in striatal GABAergic neurons. Because cortical GABAergic neurons are a minority group of neurons scattered across cortex, disappearance of *LacZ* staining in these GABAergic neurons cannot be directly visualized in the cortex of *Sp4* heterozygous mice carrying a *Dlx6a-Cre* transgene. However, *Dlx6a-Cre* expression in cortical GABAergic neurons was readily detected in the same *Dlx6a-Cre* mouse line when breeding with *ROSALacZ* (*Gtrosa26^tm1Sor^*) reporter mice (http://www.informatics.jax.org/recombinase/specificity?id=MGI:3758328&systemKey=4856356).

**Figure 1.**
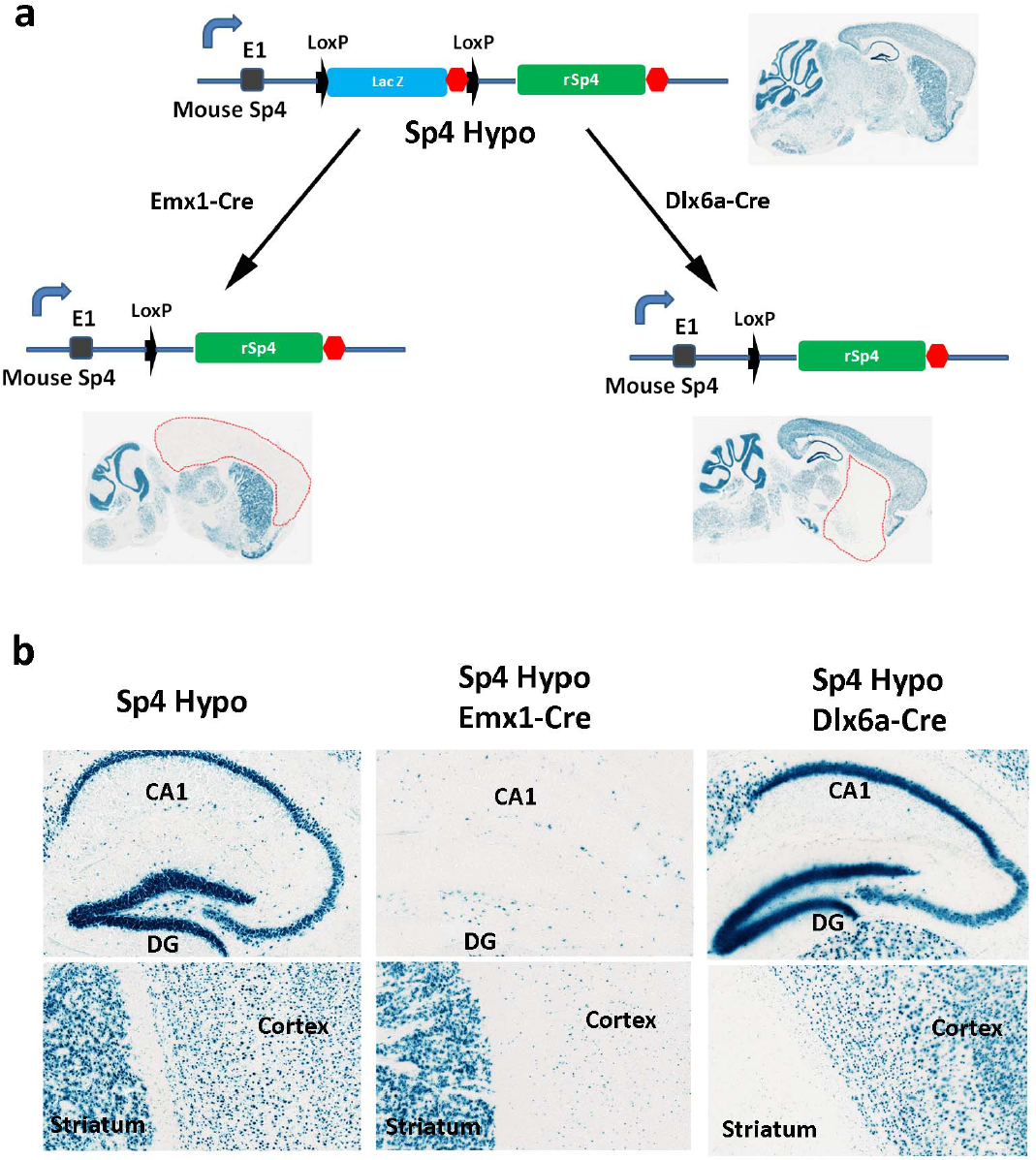
Restoration of rat *Sp4* gene in forebrain excitatory neurons and GABAergic neurons respectively. **(a)** A nuclear *LacZ* expression cassette was capped with a splicing acceptor and further flanked by two Lox P sites. The Lox P flanked *LacZ* was inserted in the first intron of mouse *Sp4* gene. Expression of *LacZ* gene revealed that the *Sp4* gene is specifically expressed in neuronal cells in mouse brain. After breeding with *Emx1-Cre* and *Dlx6a-Cre* mouse lines, the *LacZ* gene cassette was excised in forebrain excitatory neurons and GABAergic neurons respectively. The removal of the *LacZ* cassette allows expression of the downstream rat *Sp4* gene that was also capped with a splicing acceptor to splice with mouse exon 1 to generate a functional full-length Sp4 gene. The absence of blue staining in neuronal cells indicates restoration of the *Sp4* gene. Because mouse cortex predominantly consists of excitatory neurons, *LacZ* blue staining almost completely disappeared in the cortex of *Sp4* hypomorphic mice carrying the *Emx1-Cre* gene. In contrast, mouse striatum predominantly contains GABAergic neurons, the *LacZ* blue staining almost completely disappeared in the striatum of *Sp4* hypomorphic mice carrying *Dlx6a-Cre* gene. **(b)** Under higher magnification, neurons that were not derived from *Emx1* lineage remain blue in the cortex of *Sp4* hypomorphic mice carrying *Emx1-Cre* gene. Many of them are GABAergic neurons.

RT-PCR confirmed restoration of rat *Sp4* (*rSp4*, capped with a splicing acceptor) RNA expression in the cortex of (*Sp4 ^Hypo/Hypo^; Emx1^+/+(Cre)^*) rescue mice (Figure 2a). A lower level of *Sp4* RNA amplification from the striatum of the rescue mice likely comes from the restoration of *Sp4* gene in a few striatal neurons derived from *EMX1-Cre* lineage (Cocas *et al*, 2009). Western blot provided further validation of restoration of the Sp4 transcription factor in the cortex of the (*Sp4 ^Hypo/Hypo^; Emx1^+/+(Cre)^*) rescue mice (Figure 2b). Restoration of the Sp4 transcription factor was also confirmed in the striatum of (*Sp4 ^Hypo/Hypo^; Tg(Dlx6a-Cre)*) rescue mice (Figure 2c). An increased level of Sp4 protein was also detected from the cortex of GABAergic *Sp4* rescue mice (*Sp4 ^Hypo/Hypo^; Tg(Dlx6a-Cre)*), because of restoration of *Sp4* gene in cortical GABAergic neurons.

**Figure 2.**
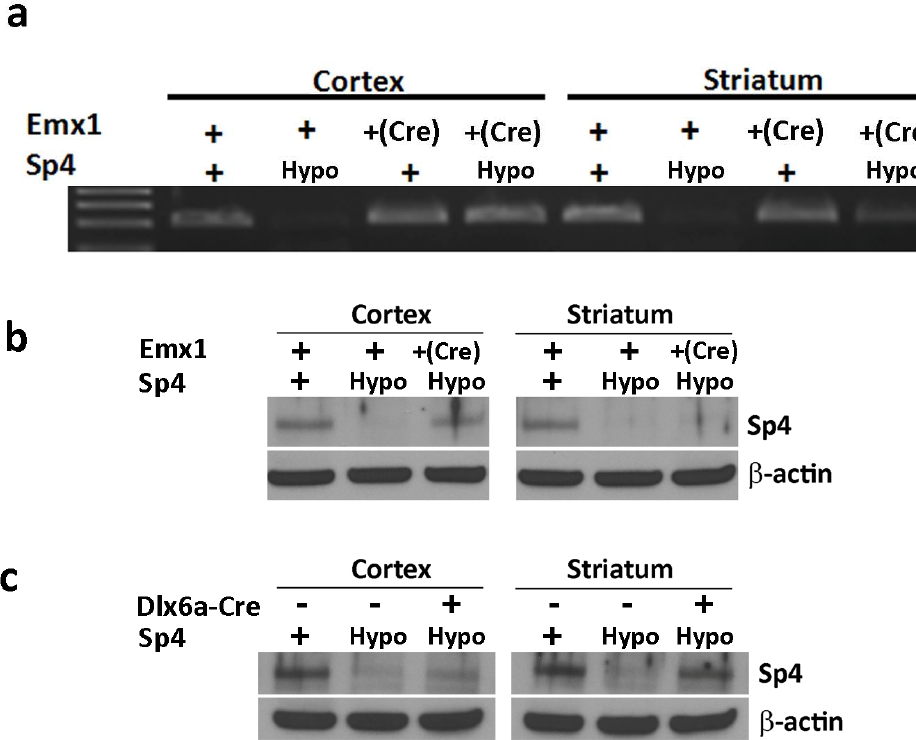
Sp4 expression in neuron-specific *Sp4* rescue mice. (a) RT-PCR was conducted to confirm rat *Sp4* RNA expression with one primer from mouse exon 1 and the other one from rat *Sp4* gene (Zhou *et al*, 2005). The presence of *Emx1-Cre* restored *Sp4* expression in the cortex of the rescue mice (*Sp4^Hypo/Hypo^; Emx1^+/+(Cre)^*). Some expression of rat Sp4 in the striatum of the rescue mice likely comes from a few *Emx1* expressing neurons. **(b)** Western blot confirmed expression of Sp4 protein in the cortex of rescue mice (*Sp4^Hypo/Hypo^; Emx1^+/+(Cre)^*). **(c)** Western blot confirmed expression of Sp4 protein in the striatum of rescue mice (*Sp4^Hypo/Hypo^; Tg(Dlx6a-Cre)*). A low level of Sp4 protein from the cortex of the rescue mice likely come from restored Sp4 expression in cortical GABAergic neurons.

## Behavioral characterization of the *Emx1-Cre* rescue mice

Deficits in PPI and fear learning as well as ketamine–induced hyperlocomotion were examined in the two different neuron-specific *Sp4* rescue mice. In the *Emx1-Cre* rescue cohort, *Sp4* hypomorphic mice showed significantly reduced PPI (F(1,80)=18.23, p < 0.0001) (Figure 3a), which was not rescued by restoration of the *Sp4* gene in forebrain excitatory neurons in (*Sp4 ^Hypo/Hypo^; Emx1 ^+/+(Cre)^*) rescue mice (*Sp4* X *Emx1-Cre* interaction, F(1,80)<1, ns). No sex effects were observed. Acquisition of fear learning was also reduced in *Sp4* hypomorphic mice, which was not rescued by introduction of the *Emx1-Cre* gene (Figures S2a to S2c). We examined the locomotor response of these mice to ketamine (50 mg/kg) using video-tracking (VT) equipment. Before ketamine treatment, there was no significant *Sp4* gene effect on distance traveled during habituation. After ketamine injection, *Sp4* hypomorphic mice exhibited X ketamine interaction (F(1,79)= 12.97, p = 0.0006) on distance traveled). No *Sp4* X *Emx1-Cre* X ketamine interaction was detected (F(1,79)<1, ns). The rescue mice (*Sp4 ^Hypo/Hypo^; Emx1 ^+/+(Cre)^*) displayed similar ketamine-induced hyperlocomotion as the (*Sp4 ^Hypo/Hypo^; Emx1^+/+^*) *Sp4* hypomorphic mice after ketamine injection, suggesting that there was no rescue of ketamine hypersensitivity.

**Figure 3.**
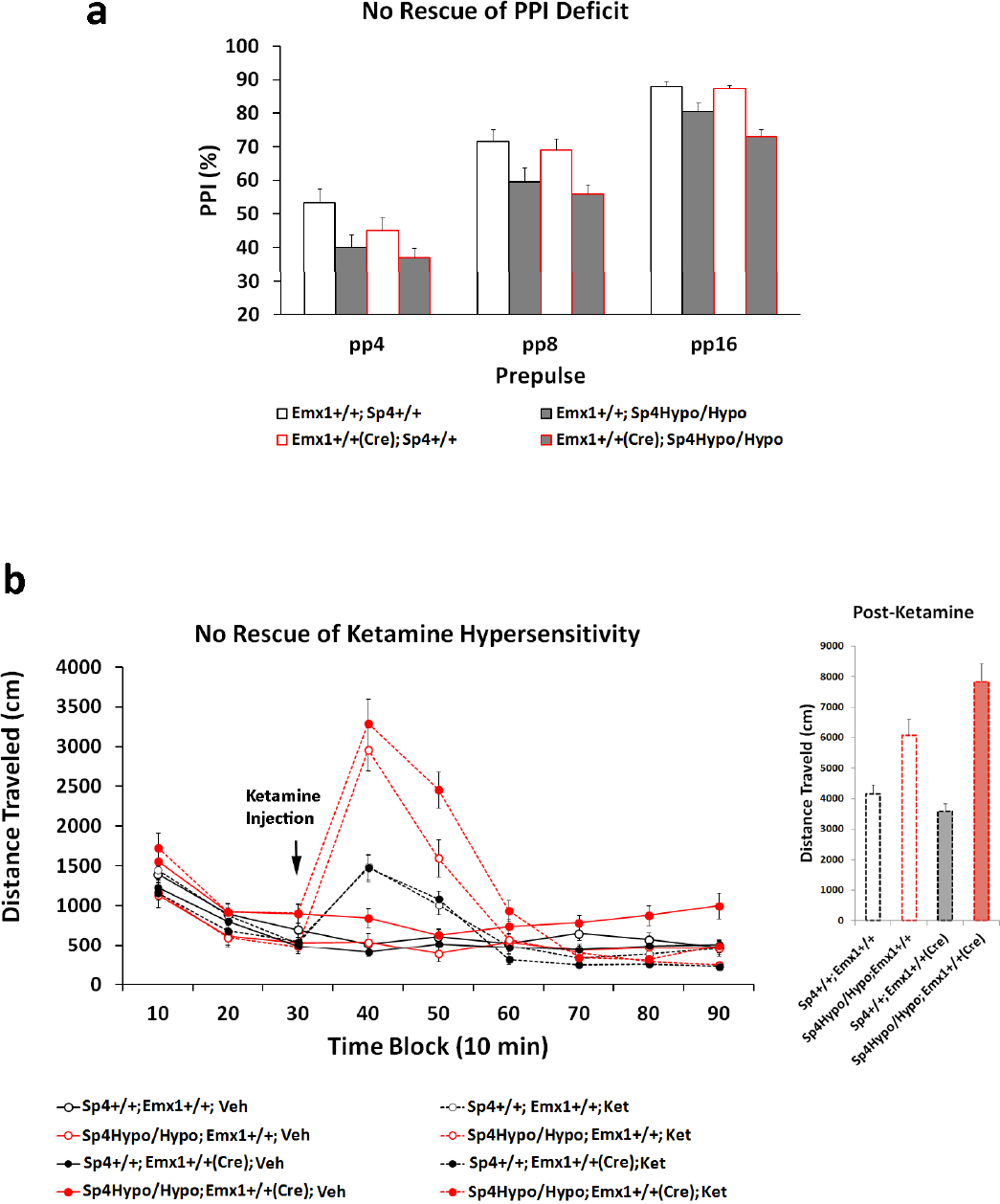
Characterization of rescue mice with restoration of *Sp4* gene only in forebrain excitatory neurons. Male and female mice were balanced in each genotype. There were 18 WT (*Sp4^+/+^; Emx1^+/+^*), 20 *Sp4* hypomorphic (*Sp4^Hypo/Hypo^; Emx1^+/+^*), 19 WT carrying *Emx1-Cre* (*Sp4^+/+^; Emx1^+/+(Cre)^*), 27 rescue mice (*Sp4^Hypo/Hypo^; Emx1^+/+(Cre)^*) for behavioral analyses. **(a)** Prepulse inhibition was conducted in these mice. No sex effect was found. *Sp4* hypomorphic mice displayed PPI deficits across three different levels of prepulse compared to wildtype mice regardless of the presence or absence of the *Emx1-Cre* gene. **(b)** All mice were habituated for 30 min before injection of ketamine (50 mg/kg). Injection of ketamine significantly increased locomotor activity in all 4 genotypes. However, there was a significantly larger increase of locomotor activity in *Sp4* hypomorphic mice with or without the *Emx1-Cre* gene than in wildtype mice. There was no interaction between *Sp4* X *Emx1-Cre* X Ketamine. Error bar: SEM

## Behavioral characterization of the *Dlx6a-Cre* rescue mice

In the *Dlx6a-Cre* rescue cohort, *Sp4* hypomorphic mice exhibited reduced PPI (F(1,85)=9.73, p = 0.002) (Figure 4a). However, no *Sp4* X *Dlx6a-Cre* interaction was detected (F(1,85)=1.38, ns) with the (*Sp4 ^Hypo/Hypo^; Tg(Dlx6a-Cre)*) rescue mice exhibiting similar PPI deficits to *Sp4* hypomorphic mice. Additionally, the deficient fear learning in *Sp4* hypomorphic mice was not rescued (Figure S2d). The response to ketamine was subsequently examined with the VT locomotion test. No significant *Sp4* effect was observed during habituation. After ketamine injection, a significant *Sp4* X ketamine interaction (F(1,85)=14.54, p < 0.001) was observed (Figure 4b). Interestingly, there was a significant *Sp4* X *Dlx6a-Cre* X ketamine interaction (F(1,85)=7.44, p = 0.008). The rescue mice (*Sp4 ^Hypo/Hypo^; Tg(Dlx6a-Cre)*) displayed the same time-course pattern of locomotive hyperactivity as the control mice (*Sp4 ^+/+^; Tg(Dlx6a-Cre)*) after ketamine injection, suggesting that restoration of *Sp4* in forebrain GABAergic neurons is sufficient to rescue ketamine–induced hyperlocomotion in *Sp4* hypomorphic mice.

**Figure 4.**
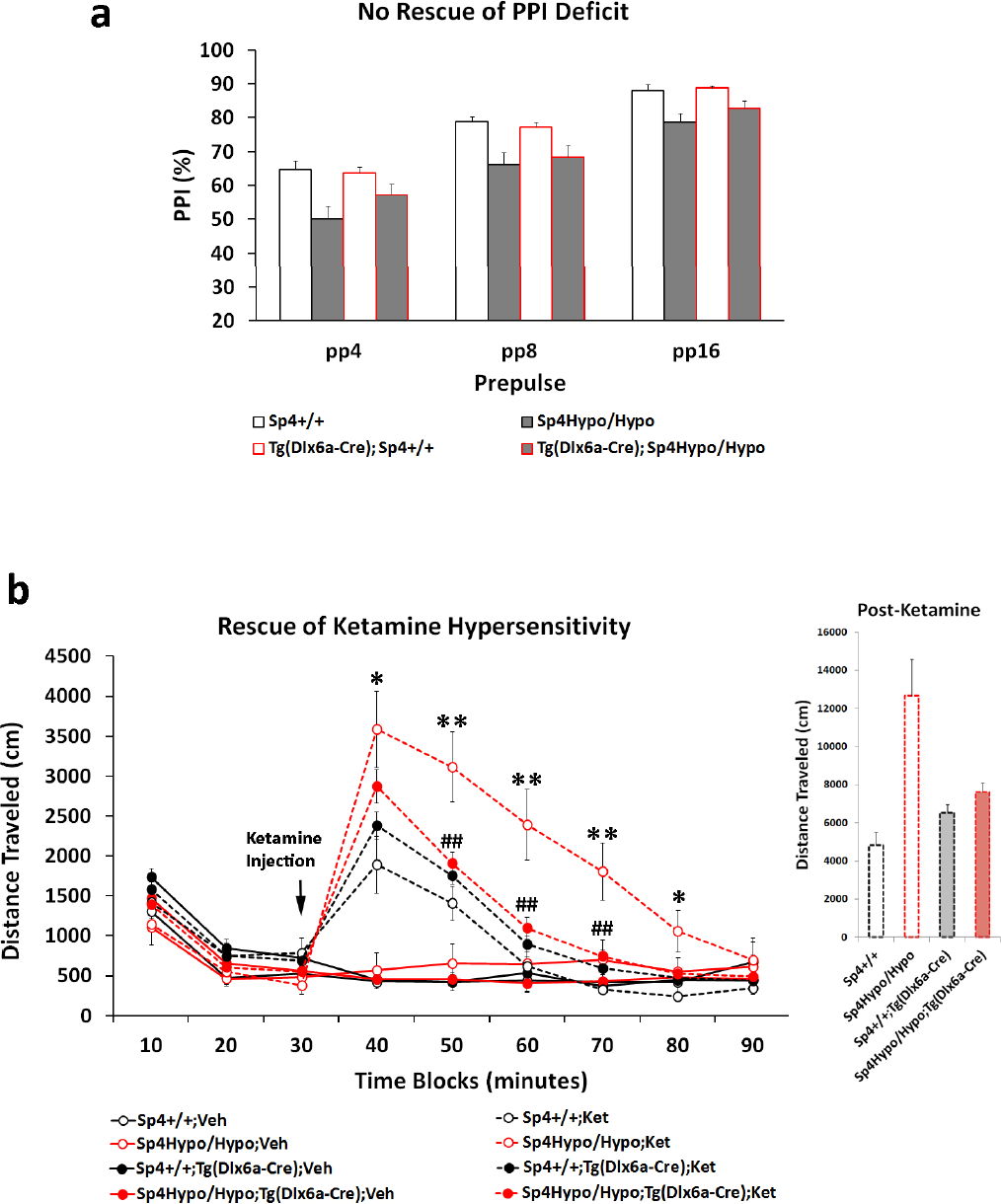
Prevention of ketamine-induced hyperlocomotion in rescue mice with restoration of *Sp4* gene only in forebrain GABAergic neurons. Male and female mice were balanced in each genotype. There were 9 WT (*Sp4^+/+^*), 14 *Sp4* hypomorphic (*Sp4^Hypo/Hypo^*), 46 WT carrying *Dlx6a-Cre* (*Sp4^+/+^; Tg(Dlx6a-Cre)*), 21 rescue mice (*Sp4^Hypo/Hypo^; Tg(Dlx6a-Cre)*) for behavioral analyses. Our breeding generated fewer wildtype and *Sp4* hypomorphic mice. More rescue mice and the control wildtype mice carrying the *Dlx6a-Cre* transgene were produced. **(a)** Prepulse inhibition was examined in all four genotypes of mice. There was no sex effect on PPI. *Sp4* hypomorphic mice displayed PPI deficits across three different levels of prepulse compared to wildtype mice. No interaction between *Sp4* X *Dlx6a-Cre* was observed. **(b)** After 30 min habituation in VT chamber, mice were injected with ketamine (50mg/kg). As expected, *Sp4* hypomorphic mice displayed significant more locomotor activity after ketamine injection (F(1,85)=14.54, p < 0.001). *Post hoc* analysis revealed more distance traveled in the *Sp4* hypomorphic mice than wildtype mice at 10 (* p < 0.05), 20 (** p < 0.01), 30 (** p < 0.01), 40 (** p < 0.01), and 50 min (* p < 0.05) post-ketamine injection. A significant *Sp4* X *Dlx6a-Cre* X ketamine interaction was found (F(1,85)=7.44, p = 0.008). *Post hoc* analysis revealed significant differences in distance traveled between the *Sp4* hypomorphic mice in the absence or presence (rescue) of the *Dlx6a-Cre* transgene at 20 (^##^ p < 0.01), 30 (^##^ p < 0.01), 40 (^##^ p < 0.01) min post-ketamine injection. The rescue mice (*Sp4 ^Hypo/Hypo^*; *Tg(Dlx6a-Cre*) displayed no difference in locomotor activity from the wildtype mice carrying the *Dlx6a-Cre* gene, suggesting an almost complete rescue of the excessive response to ketamine. Error bar: SEM.

## Discussion

Genetic manipulations are powerful tools for disentangling the complex phenotypes. Conventional and conditional gene-knockout strategies can determine whether a gene is necessary for a particular phenotype in a specific type of cells. Conditional rescue of a phenotype in knockout mice via restoration of the gene in only one specific cell type can determine whether the gene in that specific cell type is necessary and sufficient for controlling the phenotype. Many behaviors are complex and likely modulated by interactions between different types of neurons from different brain regions. Given that different types of neurons are involved in the modulation of behavior, it is not surprising that *Sp4* gene restoration in only one specific type of neurons may not be sufficient to rescue behavioral phenotypes. Indeed, we failed to rescue deficient PPI and fear learning in *Sp4* hypomorphic mice regardless of whether *Sp4* gene was restored in forebrain excitatory neurons or GABAergic neurons. Importantly however, we found that restoration of the *Sp4* gene in GABAergic neurons was sufficient to rescue ketamine-induced hyperlocomotion in the *Sp4* hypomorphic mice.

*Sp4* hypomorphic mice (*Sp4 ^Hypo/Hypo^*) from the *Dlx6a-Cre* rescue cohort displayed a prolonged response to ketamine than the wiltype control mice. However, *Sp4* hypomorphic mice (*Sp4 ^Hypo/Hypo^; Emx1^+/+^*) from the *Emx1-Cre* rescue cohort had a similar time-course response to ketamine with their sibling wiltype mice. The prolonged response to ketamine in the *Sp4* hypomorphic mice from the *Dlx6a-Cre* rescue cohort is a more typical response of *Sp4* hypomorphic mice according to our previous studies (Ji *et al*, 2013). Why did the same dosage of ketamine not elicit a prolonged response to ketamine in the *Sp4* hypomorphic mice from the *Emx1-Cre* rescue cohort? We speculate that there may be still subtle differences between the genetic backgrounds of the *Emx1-Cre* and *Dlx6a-Cre* mouse lines despite the fact that both *Emx1-Cre* and *Dlx6a-Cre* mouse lines were backcrossed for more than 6 generations. The genetic background of the *Emx1-Cre* rescue cohort may have been overall less sensitive to ketamine than that of the *Dlx6a-Cre* rescue cohort. We consider that different time-course responses to ketamine may be generated by differential ketamine sensitivities between mouse cohorts or different dosages of ketamine. Indeed, increasing dosages of ketamine produced more locomotor activity and prolonged responses (Irifune *et al*, 1991). This observation may explain why the same dosage of ketamine generated different time-courses in the locomotor responses between the two different mouse cohorts.

Ketamine acts primarily, although not exclusively (Kapur and Seeman, 2002), as a noncompetitive antagonist of NMDAR receptors that are present in both excitatory and GABAergic neurons in different brain regions. Both cortical GABAergic inhibitory and excitatory pyramidal neurons have been suggested to be the primary targets of NMDAR antagonists (Homayoun and Moghaddam, 2007; Moghaddam *et al*, 1997; Rotaru *et al*, 2011). Our studies suggest that the mouse *Sp4* gene is essential for forebrain GABAergic neurons to dampen locomotive responses to ketamine. Reduced *Sp4* expression may impair the ability of GABAergic neurons to control locomotor hyperactivity in *Sp4* hypomorphic mice after ketamine injection (Ji *et al*, 2013). In contrast, the *Sp4* gene expression appears dispensable in excitatory neurons in controlling locomotor response to ketamine. Since the genetic restoration of the *Sp4* gene in *Dlx6a-Cre* lineage cells starts from early embryogenesis, our studies cannot differentiate whether altered development (if there is) and/or function of forebrain GABAergic neurons are responsible for ketamine hypersensitivity in *Sp4* hypomorphic mice. Although we did not find gross abnormalities in the brains of *Sp4* hypomorphic mice (Ji *et al*, 2013; Zhou *et al*, 2005), we cannot rule out subtle structural alterations in GABAergic neurons. As for molecular mechanisms, NMDAR has been proposed as the primary target of ketamine. Nmdar1 protein, but not mRNA, was down-regulated in the brains of *Sp4* hypomorphic mice (Sun *et al*, 2015; Zhou *et al*, 2010). Those findings however, came from mouse brain tissue that contained different types of neurons and glial cells, and hence do not necessarily exclude the *Nmdr1* gene as a direct target of the Sp4 transcription factor in a small group of specific neurons. There is still a possibility that the Sp4 transcription factor may directly regulate mRNA expression of *Nmdar1* gene in some forebrain GABAergic neurons (Krainc *et al*, 1998). Forebrain GABAergic neurons consist of striatal neurons and different cortical GABAergic neurons, it remains to be investigated which subset of GABAergic neurons is responsible for the rescue effects. Investigation of molecular mechanisms can eventually be conducted after identification of the critical GABAergic neurons. It should be kept in mind that other receptors and channels in addition to Nmdar receptors may also mediate the effects of ketamine in *Sp4* hypomorphic mice (Jamie Sleigha, 2014). In the future, restoration of the *Sp4* gene in forebrain GABAergic neurons via temporal and regional-specific (e.g. cortex or striatum) control of Cre-mediated DNA recombination may clarify the role of development of GABAergic neurons and functions of regional GABAergic neurons in controlling locomotor responses to ketamine.

Abnormalities of cortical GABAergic neurons have been reported consistently in the postmortem brains from patients with schizophrenia (Lewis *et al*, 2005). Ketamine produces schizophrenia-like behavioral phenotypes in healthy people and exacerbates symptoms in schizophrenia patients. Our studies suggest that *SP4*, deleted in some patients with schizophrenia, could be a missing piece that links a genetic susceptibility gene with ketamine hypersensitivity and abnormal function of GABAergic neurons in schizophrenia. The influence of the *Sp4*-GABAergic-ketamine pathway may not be limited to the regulation of locomotor activity in response to ketamine. Given that ketamine functions as a potent antidepressant, it will be interesting to investigate whether the *Sp4*-GABAergic-ketamine pathway may also be involved in the regulation of depressive behaviors. Indeed, our most recent studies suggested that *Sp4* hypomorphic mice displayed depressive-like behaviors (Young et al., submitted). In the future, such depressive-like phenotypes will be examined after ketamine treatment in *Sp4* hypomorphic mice with restoration of the *Sp4* gene in GABAergic neurons.

## Acknowledgements

This study is supported by R01MH073991 (X.Z.) and the Veterans Affairs VISN 22 Mental Illness Research, Education, and Clinical Center.

